# Plcl1 Regulates Hematopoietic Stem Cell Function During Aging and Stress by Modulating Calcium Dynamics

**DOI:** 10.1101/2025.09.15.674672

**Authors:** Tomohiro Yabushita, Yosuke Tanaka, Tsuyoshi Fukushima, Kanako Wakahashi, Terumasa Umemoto, Hitoshi Takizawa, Susumu Goyama, Toshio Kitamura, Akira Nishiyama, Tomohiko Tamura, Satoshi Yamazaki, Toshio Suda

## Abstract

Long-term hematopoietic stem cells (HSCs) can generate all blood lineages but typically remain quiescent, becoming activated only in response to acute stress. We previously demonstrated that quiescent HSCs exhibit heterogeneity in intracellular calcium levels. However, the mechanisms underlying this heterogeneity and its physiological relevance remain unclear. Herein, we identify phospholipase C-like 1 (*Plcl1*), a noncatalytic protein that binds inositol 1,4,5-trisphosphate (IP_3_), as being selectively enriched in the most quiescent HSC subset. Loss-of-function studies revealed that *Plcl1* deficiency at steady state reduced basal intracellular calcium levels and skewed the HSC compartment toward CD41⁺ subsets while preserving overall HSC numbers and long-term reconstitution capacity. Under acute hematopoietic stress, *Plcl1* loss accelerated and amplified platelet rebound and the expansion of non-canonical megakaryocyte progenitors (ncMkPs), indicating activation of the thrombopoietic bypass pathway. In aged HSCs, *Plcl1* deficiency exacerbated aging-related features, including expansion of the HSC pool, accumulation of CD41⁺ HSCs and ncMkPs, and myeloid-skewed differentiation with impaired competitive reconstitution. These changes were accompanied by diminished induction of calcium-responsive immediate-early genes. Collectively, we identified *Plcl1* as an intrinsic regulator that stabilizes calcium dynamics in HSCs, thereby restraining stress- and aging-associated megakaryocytic priming and preserving stem cell function.

## Introduction

Hematopoietic stem cells (HSCs) sustain lifelong hematopoiesis by leveraging their ability to self-renew and differentiate into multiple lineages. A hallmark of long-term HSCs (LT-HSCs) is their quiescent state, defined by arrest in the G_0_ phase of the cell cycle, which preserves genomic integrity, prevents replicative exhaustion, and maintains their long-term regenerative capacity^1,2^. This dormant state, in turn, is actively sustained by complex transcriptional, epigenetic, and metabolic programs and extrinsic niche-derived factors^3–7^. However, the complete set of molecular pathways governing the G_0_ state in HSCs remains unclear.

Previous studies have shown that LT-HSCs exhibit substantial functional heterogeneity^8,9^. Recently, calcium (Ca^2+^) signaling has emerged as a key determinant of HSC fate^10,11^. Cytokines such as stem cell factor (SCF) and thrombopoietin (TPO) increase intracellular Ca^2+^ levels, promoting HSC activation and differentiation^10^. Consistent with their quiescent and undifferentiated state, LT-HSCs maintain substantially lower basal Ca^2+^ levels than lineage-committed progenitors^10,12^. However, in our previous study, when focusing on the quiescent LT-HSC compartment defined by G_0_ markers, HSCs with relatively elevated intracellular Ca^2+^ levels exhibited enhanced dormancy, bone marrow reconstitution, and self-renewal capacity^13^. Taken together, these findings indicate that intracellular Ca^2+^ dynamics play multifaceted roles in HSC biology and highlight the importance of elucidating how intracellular Ca^2+^ levels are tightly regulated in HSCs and how their disruption affects stem cell function.

Recent evidence implicates dysregulated intracellular Ca^2+^ dynamics as a major contributor to HSC aging^14^. During the physiological HSC aging, abnormal intracellular Ca^2+^ enrichment promotes mitochondrial dysfunction, impairs regenerative capacity, and intensifies myeloid-biased differentiation in HSCs^14^. Another hallmark of aged HSCs is the expansion of megakaryocyte-biased subsets, which preferentially support thrombopoiesis via a non-canonical bypass pathway^15–18^. Beyond the classical route through multipotent progenitors (MPPs) and canonical CD48⁺ megakaryocyte progenitors (MkPs), lineage-tracing and single-cell analyses have revealed that HSCs can directly generate CD48^−^/low non-canonical MkPs, accelerating platelet production^19–21^. This bypass pathway is likewise engaged during acute stress hematopoiesis, such as 5-fluorouracil (5-FU) or poly I:C, enabling rapid platelet replenishment^21,22^. Although such acute response is essential for short-term recovery, repeated or chronic stress accelerates HSC functional decline and worsens age-associated lineage bias^22,23^.

Phospholipase C-like 1 (*Plcl1*) is a non-catalytic member of the phospholipase C family that retains an inositol 1,4,5-trisphosphate (IP_3_)-binding domain but lacks enzymatic activity^24,25^. In non-hematopoietic tissues, *Plcl1* has been demonstrated to fine-tune intracellular Ca^2+^ dynamics by modulating IP_3_ receptor signaling. When IP_3_ concentrations are elevated, Plcl1 can attenuate receptor-mediated Ca^2+^ release by competing for IP_3_ binding; under physiological conditions, however, it stabilizes IP_3_ and facilitates efficient receptor signaling^26–28^. Despite these insights, the functional relevance of Plcl1 in HSC regulation remains uncharacterized. In this study, we examine Plcl1 as an intrinsic regulator of Ca^2+^ homeostasis in HSCs by integrating transcriptomic analyses, genetic loss-of-function models, and transplantation assays to delineate its role in maintaining quiescence, balanced lineage output, and stress responses during aging and acute hematopoietic stress.

## Materials and Methods

### Mice

The *Plcl1* knockout (*Plcl1*-KO) mouse line, also known as the *Prip1*-deficient model, was first established and characterized by Dr. Kanematsu and Dr. Hirata^29^. To assess HSC quiescence in vivo, we used *Vav1*-Cre;*Rosa^R26R-mVenus-p27K-/R26R-wt^*mice, wherein quiescent hematopoietic cells are visualized by the expression of mVenus–p27K^−^, a fluorescent reporter that does not bind Cdks and thereby specifically marks G_0_-phase cells without altering cell cycle dynamics^13^. All experiments were conducted using male mice aged 12–16 weeks. For aging studies, male mice aged 18–21 months were used. Acute stress hematopoiesis was induced by intraperitoneal injection of 150 mg/kg 5-FU (Kyowa Kirin) or by intravenous injection of 2 μg anti-mouse CD42b antibody (R300, Emfret Analytics). All animals were maintained under specific pathogen-free conditions at the Center for Animal Resources and Development (CARD), Kumamoto University. All procedures were approved by the Animal Care and Use Committee of Kumamoto University.

### Cell preparation

Bone marrow (BM) cells were isolated by gently crushing the pelvis, femurs, and tibias in Dulbecco’s Modified Eagle Medium (Sigma) supplemented with 10% fetal bovine serum (FBS; Biowest). Following red blood cell lysis with ACK buffer (Thermo Fisher Scientific), the suspensions were filtered through 100 µm strainers and washed once with Dulbecco’s phosphate-buffered saline (PBS; Sigma) containing 2% FBS. Nucleated cells were subsequently counted using Turk’s solution. For HSC enrichment, the BM cells were magnetically labeled with microbead-conjugated anti–c-Kit antibodies (Miltenyi Biotec), fractionated using the autoMACS Pro Separator (Miltenyi Biotec), and stained with fluorescence-conjugated antibodies for analysis and sorting.

### BM and peripheral blood analysis

Fluorescence-activated cell sorting (FACS) staining was performed using the following antibodies: anti-c-kit (2B8), anti-CD150 (TC15-12F12.2), anti-CD48 (HM48-1), anti-EPCR (eBio1560), anti-CD41 (MWReg30), anti-Sca-1 (D7), anti-Ter-119 (TER-119), anti-CD4 (GK1.5), anti-CD8a (53-6.7), anti-B220 (RA3-6B2), anti-Gr-1 (RB6-8C5), and anti-Mac-1 (M1/70). All antibodies were purchased from BioLegend unless stated otherwise. Flow cytometric analysis and cell sorting were conducted using the FACSAria III instrument (BD Biosciences). Data were analyzed using FlowJo software, version 10.9.0 (Beckman Coulter). Cell populations were defined as follows: KSL (Lineage^−^c-Kit^+^Sca-1^+^), MPP2 (CD150^+^CD48^+^ KSL), MPP3/4 (CD150^−^CD48^+^ KSL), ST-HSC (CD150^−^CD48^−^ KSL), LT-HSC (HSC; CD150^+^CD48^−^ KSL), LK (Lineage^−^c-Kit^+^Sca-1^−^), MkP (CD150^+^CD41^+^LK), cMkP (canonical MkP; CD48^+^MkP) and ncMkP (non-canonical MkP; CD48^−/low^ MkP). Following 5-FU treatment, HSCs were defined as Lin⁻EPCR⁺CD150⁺CD48^−^ cells (L^−^ESLAM), based on previously validated criteria that reliably identify functionally pure HSCs under both steady-state and stress conditions^30,31^.

### Cell cycle analysis

The cell cycle status of HSCs and haematopoietic stem and progenitor cells (HSPCs) was assessed using the PerFix EXPOSE Kit (Phospho-Epitopes Exposure Kit; Thermo Fisher Scientific) in accordance with the manufacturer’s protocol. Freshly isolated BM cells were surface-stained with lineage and HSC markers, following which they were fixed and permeabilized using the kit’s reagents. Intracellular Ki-67 was detected using Alexa-Fluor-647-conjugated anti–Ki-67 antibody (Ki-67, BioLegend), and DNA content was assessed by staining with propidium iodide (BioLegend). Samples were analyzed by flow cytometry as described above.

### Transplantation

For competitive transplantation, FACS-purified LT-HSCs (CD150⁺CD48⁻ KSL) from wild-type (WT) or *Plcl1*-KO (CD45.2) mice were intravenously co-injected with 2 × 10^6^ unfractionated CD45.1^+^ competitor BM cells into lethally irradiated CD45.1^+^ recipients (9.5 Gy), using 100 LT-HSCs for young mice and 250 for aged mice. Donor chimerism and lineage contribution were assessed in peripheral blood every 4 weeks up to 16 weeks using antibodies against CD45.1 (A20), CD45.2 (104), Gr-1(RB6-8C5), CD11b (M1/70), B220 (RA3-6B2), CD4(GK1.5), and CD8a(53-6.7). These antibodies were purchased from BioLegend.

### Intracellular calcium analysis

Freshly isolated HSCs were incubated with 0.5 μM Fluo-4 AM and 0.04% Pluronic F-127 (both from Thermo Fisher Scientific) for 1 h at 37 °C in 5% CO_2_. Following staining, the cells were washed with PBS containing 2% FBS and resuspended in the same buffer for analysis. Fluorescence was measured using the FITC channel on a BD FACSAria III cell sorter, and flow cytometric data were analyzed using FlowJo software (BD Biosciences).

### RNA extraction and quantitative polymerase chain reaction (qPCR)

Total RNA was extracted from LT-HSCs using the RNeasy Micro Kit (QIAGEN). First-strand cDNA was synthesized using the ReverTra Ace™ qPCR RT Kit (Toyobo) according to the manufacturer’s instructions. qPCR was performed on a Roche LightCycler96 system using the Luna Universal qPCR Master Mix (New England Biolabs). Gene expression levels were normalized to *Gapdh*. The primer sequences were as follows: *Plcl1* forward 5′-AGAAGAAGATTAGCAGTGCCCA-3′ and reverse 5′-TCGGTTGTAGATGCGAGAGTT-3′; *Gapdh* forward 5′-CATCACTGCCACCCAGAAGACTG-3′ and reverse 5′-ATGCCAGTGAGCTTCCCGTTCAG-3′.

### RNA-seq

As previously described^10^, 100 FACS-purified LT-HSCs were used for RNA-seq library construction. First-strand cDNA synthesis was performed using the PrimeScript RT reagent kit (Takara) and not-so-random primers. Second-strand synthesis was carried out using Klenow Fragment (3′–5′ exo⁻; New England Biolabs) and complementary primers. Following purification, the resulting double-stranded cDNA was used for library preparation and amplification with the Nextera XT DNA Sample Preparation Kit (Illumina), according to the manufacturer’s protocol. Sequencing was conducted on an Illumina NextSeq 500 platform to generate 75-bp single-end reads. Subsequently, FASTQ files were uploaded to the Galaxy platform (https://usegalaxy.org) for downstream analysis. Sequencing reads were aligned to the mouse reference genome (mm39) using HISAT2, and gene-level counts were generated with FeatureCounts. These count data were then analyzed using the RNAseqChef web-based transcriptome platform^32^. Enrichment analysis was conducted in RNAseqChef using the Pathway Interaction Database and Gene Ontology (GO) Biological Process categories to identify pathways associated with differentially expressed genes. Moreover, previously generated RNA-seq datasets (GEO: GSE139013 and GSE138884) were reanalyzed to compare gene expression profiles across four HSC fractions: G_0_M-negative HSCs, G_0_M-low HSCs, G_0_M-high & Ca^2+^-low HSCs, and G_0_M-high & Ca^2+^-high HSCs. Complementary gene expression data were also obtained from the Blood Stem Cell Atlas^33^ and Gene Expression Commons^34^ for broader validation.

### Statistical analysis

All statistical analyses were performed using GraphPad Prism software. Data are presented as mean ± standard deviation (SD) or standard error of the mean (SEM), as indicated. Comparisons between two groups were performed using unpaired two-tailed Student’s t-tests. For comparisons involving multiple groups, one-way or two-way analysis of variance followed by Bonferroni post hoc testing was conducted. A *p*-value < 0.05 was considered statistically significant.

## Results

### *Plcl1* Expression Characterizes the Most Functionally Competent HSC Subset

To identify novel regulators associated with HSC quiescence and functional capacity, we reanalyzed our previously published transcriptomic dataset obtained from G_0_ marker (G_0_M) reporter mice (*Vav1-Cre;Rosa^R26R-G0M/R26R-wt^*mice). In this model, G_0_-phase HSCs were further subdivided according to intracellular Ca^2+^ levels using CaSiR, a cell-permeant fluorescent Ca^2+^ indicator^13^. This analysis identified *Plcl1* as one of the top differentially expressed genes in the G_0_M-high/Ca^2+^-high HSC subset, which corresponds to the most quiescent and functionally potent population **(Figure 1A)**. To validate this observation, we analyzed a publicly available single-cell RNA-seq dataset of adult murine hematopoiesis^33^. *Plcl1* expression was significantly higher in LT-HSCs (Lin⁻Sca-1⁺c-Kit⁺CD34⁻Flk2⁻) than in multipotent and lineage-committed progenitors **(Figure S1A–B)**. Single-cell expression analysis also demonstrated a strong positive correlation with canonical HSC markers (CD150, EPCR, and Sca-1) and an inverse correlation with Flk2, a marker of progenitor commitment **(Figure S1C)**. We next queried Gene Expression Commons^34^, a platform that assigns binary activity states (“active” or “inactive”) across diverse hematopoietic populations. This approach identified an HSC-specific gene set (active in HSCs but inactive in other HSPCs and differentiated lineages), comprising 117 candidates, including *Hoxb5* and *Tcf15* (**Figure S1D–E)**. To further enrich for the most quiescent population, we compared vWF^+^ and vWF⁻ HSCs and identified 14 vWF⁺-HSC-specific genes. Notably, *Plcl1* was among these vWF⁺-HSC-specific genes (**Figure 1B**, **Figure S1E**). Finally, consistent with its association with stem cell dormancy, *Plcl1* expression was rapidly downregulated upon *in vitro* culture of HSCs, suggesting that its expression is closely linked to the maintenance of stem cell quiescence **(Figure S1F)**.

**Figure 1.**
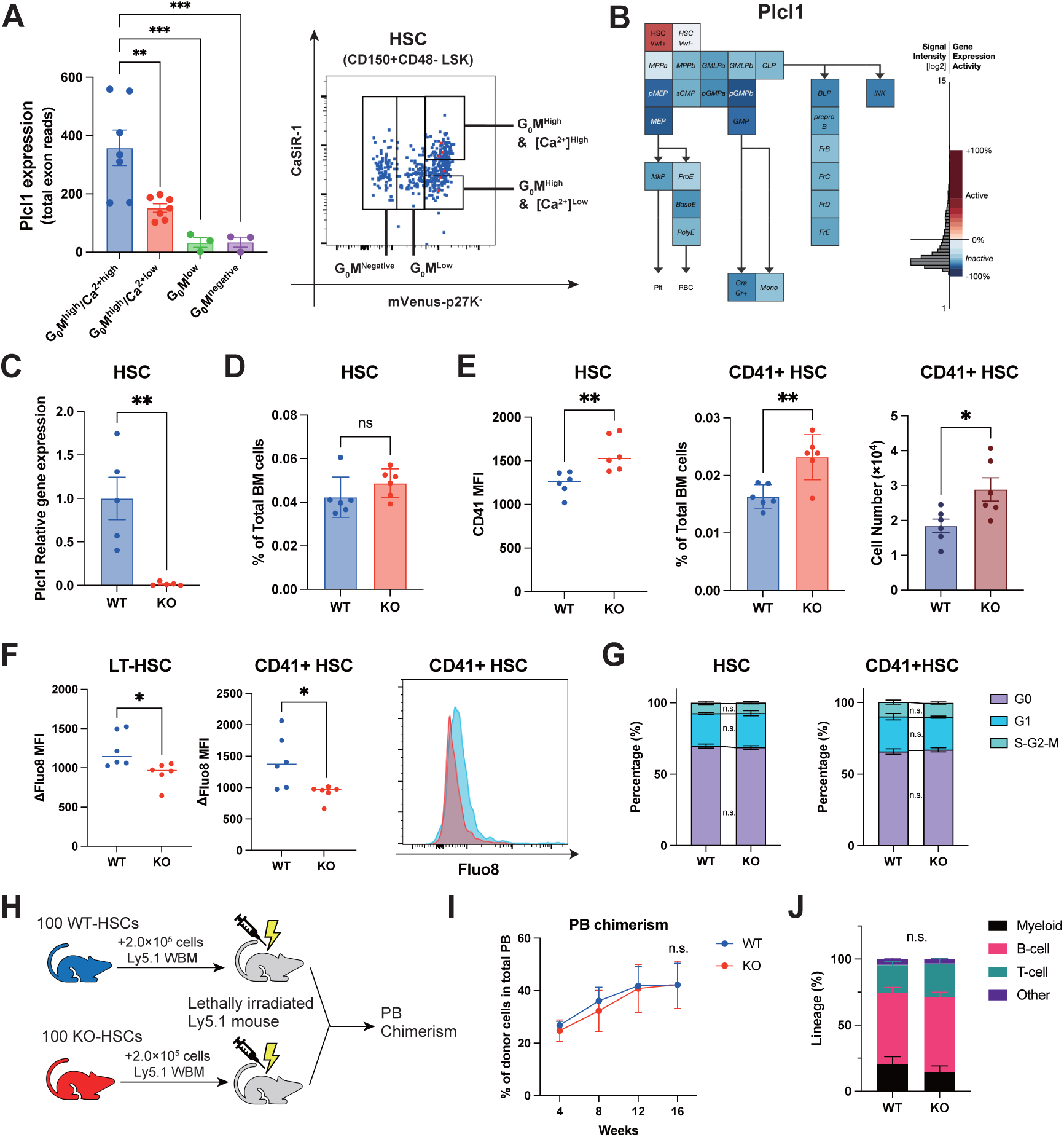
*Plcl1* is enriched in quiescent HSCs and regulates their Ca^2+^ dynamics and subset composition. **(A)** Quantitative comparison of *Plcl1* expression across HSC subsets gated by mVenus–p27K⁻ intensity (a G_0_ marker consisting of mVenus fused to a CDK-binding–defective p27)^54^ and CaSiR fluorescence (intracellular Ca**^2+^** indicator). \*\**p* < 0.01; \*\*\**p* < 0.001. The gating strategy shown on the right was adapted from our previous study. **(B)** Heatmap from the Gene Expression Commons database showing that *Plcl1* expression is highest in vWF⁺ HSCs (see Supplementary Fig. 1E for population abbreviations). **(C)** Quantification of *Plcl1* expression in SLAM-KSL HSCs from WT and *Plcl1*-deficient (KO) mice, assessed by qPCR (n = 5). **p < 0.01. **(D)** Frequency of LT-HSCs as a percentage of total BM cells in WT and *Plcl1*-KO mice (n = 6). ns: not significant. **(E)** CD41 mean fluorescence intensity (MFI) (left), frequency (% of total BM cells, middle), and absolute number (right) of CD41⁺ HSCs (n = 6). *p < 0.05, **p < 0.01. **(F)** Intracellular Ca^2+^ levels measured by Fluo-8 in HSCs (left), CD41⁺ HSCs (middle), and CD41⁺ HSCs displayed in a representative histogram (right) (n = 6). *p < 0.05; ns, not significant. **(G)** Cell cycle states of HSCs (left) and CD41⁺ HSCs (right) in WT and *Plcl1*-KO mice (n = 6). ns, not significant. **(H)** Schematic overview of the competitive BM transplantation experiment. CD45.2⁺ donor cells from WT or *Plcl1*-KO mice were mixed with CD45.1⁺ competitor cells and transplanted into lethally irradiated Ly5.1 recipient mice. **(I)** Donor-derived peripheral blood chimerism at 4, 8, 12, and 16 weeks after competitive transplantation of WT or *Plcl1*-KO HSCs (n = 5). ns, not significant. **(J)** Donor-derived peripheral blood lineage distribution at 16 weeks post-transplantation in recipients of WT or *Plcl1*-KO HSCs (n = 5). ns, not significant.

### *Plcl1* Deficiency Shifts the HSC Compartment Toward a CD41⁺ Bias and Reduces Intracellular Ca^2+^ Levels under Steady-State Conditions

To explore the physiological role of *Plcl1* in maintaining HSC homeostasis, we analyzed *Plcl1*-deficient (*Plcl1*-KO) mice under steady-state conditions. We confirmed that *Plcl1* was expressed in LT-HSCs from WT mice but absent in *Plcl1*-KO mice **(Figure 1C)**. Peripheral blood counts, including white blood cells, hemoglobin, and platelets, were comparable between genotypes **(Figure S2A)**. Likewise, both the frequency and absolute number of HSCs remained unchanged (**Figure 1D**). In contrast, *Plcl1*-KO mice exhibited a significant increase in CD41⁺ HSCs, a subset associated with megakaryocytic lineage bias,^35–37^ while EPCR⁺ HSCs, which are linked to deeper quiescence, remained unchanged (**Figure 1E, Figure S2B**). To determine whether these phenotypic changes reflected altered intracellular signaling, we measured intracellular Ca^2+^ levels in HSCs using Fluo-8 fluorescence. Intracellular Ca^2+^ levels were significantly reduced in *Plcl1*-KO HSCs, including within the CD41⁺ subset (Figure 1F). This finding is consistent with the proposed biphasic role of *Plcl1* in IP_3_-dependent Ca^2+^ regulation, where its absence impairs IP_3_ stabilization and reduces Ca^2+^ levels under steady-state conditions^26^. Subsequently, we assessed downstream megakaryocytic commitment. Although the total number of MkPs was unchanged, CD41 expression was significantly elevated in ncMkPs (CD48^−/low^)^19–21^ from *Plcl1*-KO mice, while cMkPs (CD48⁺)^19–21^ remained unaffected **(Figure S2C–D)**. Other HSPC populations, including short-term HSCs (ST-HSCs) and MPPs, displayed no differences in abundance or intracellular Ca^2+^ levels **(Figures S2D–E)**. Cell cycle analysis revealed no differences in the proportion of G_0_-phase cells within LT-HSCs or CD41⁺ HSCs (**Figure 1G**), although a modest decrease in quiescence was observed in MPP2 cells (**Figure S2F**). To assess long-term functional capacity, we performed competitive BM transplantation by co-injecting CD45.2⁺ WT or *Plcl1*-KO donor cells with CD45.1⁺ competitor cells into lethally irradiated recipients (**Figure 1H**). Over a 16-week period, donor chimerism and multilineage reconstitution, including B cells, T cells, and myeloid cells, were comparable between the WT and *Plcl1*-KO groups (**Figure 1I–J and Figure S2G**), indicating preserved self-renewal and multilineage output in the absence of *Plcl1*. Collectively, these results demonstrate that *Plcl1* is dispensable for overall HSC maintenance under homeostatic conditions but plays a selective role in stabilizing intracellular Ca^2+^ levels and limiting the expansion of megakaryocyte-biased HSC subsets.

### *Plcl1* Attenuates Emergency Thrombopoiesis During Acute Hematopoietic Stress

Given its role in regulating intracellular Ca^2+^ and limiting megakaryocyte-biased HSC expansion under steady-state conditions, we next examined how *Plcl1* influences hematopoietic output under regenerative stress. Acute hematopoietic stress promotes platelet-biased hematopoiesis by activating a non-canonical thrombopoietic bypass pathway that originates directly from megakaryocyte-biased HSCs^21,22^. To model this response, we administered 5-FU, which induces acute hematopoietic stress followed by a pronounced platelet rebound^38^. In WT mice, *Plcl1* expression in HSCs was markedly downregulated after 5-FU administration (**Figure 2A**), suggesting a suppressive effect of *Plcl1* in stress-induced thrombopoiesis. *Plcl1*-KO mice displayed an exaggerated platelet rebound, marked by accelerated recovery, higher peak platelet counts, and extended rebound duration (**Figure 2B–C**). Recovery of other hematopoietic lineages, including erythroid and myeloid cells, was comparable between genotypes (**Figure S3A**), indicating a thrombopoiesis-specific effect. On day 6 after 5-FU administration, when HSCs typically exhibit a transient calcium surge, intracellular Ca^2+^ levels were significantly elevated in *Plcl1*-KO HSCs, particularly in the CD41⁺ subset, despite similar total HSC counts (**Figure 2D, Figure S3B–C**). By day 12, intracellular Ca^2+^ levels had normalized; nevertheless, CD41⁺ HSCs and ncMkPs remained expanded in *Plcl1*-KO mice (**Figure 2E, Figure S3D–E**), consistent with the sustained activation of the non-canonical thrombopoietic bypass pathway. To investigate the underlying molecular mechanisms, we performed RNA-seq on Lineage⁻ EPCR⁺ CD150⁺ CD48⁻ (L^-^ESLAM) HSCs isolated on days 0, 5, and 10 following 5-FU treatment. This population was selected because conventional markers such as Sca-1 and c-Kit are dynamically regulated during stress, while L^-^ESLAM HSCs provide a stable definition of LT-HSCs^30,31^. Although principal component analysis (PCA) and clustering analyses revealed substantial temporal variation in gene expression (**Figure S3F–G**), transcriptional differences between WT and *Plcl1*-KO mice remained minimal (**Figure S3H**). These findings suggest that the enhanced platelet output observed in *Plcl1*-KO mice may arise from non-transcriptional mechanisms, such as altered Ca^2+^ dynamics or post-transcriptional regulation, rather than from global transcriptional rewiring. To further validate these observations, we employed an anti-CD42b-antibody-induced immune thrombocytopenia model, which similarly engages the non-canonical bypass pathway^39^. Consistent with the 5-FU model, *Plcl1*-KO mice exhibited accelerated and sustained platelet rebound accompanied by the expansion of CD41⁺ HSCs and ncMkPs (**Figure 2F–G and Figure S3I**). Collectively, these results identify *Plcl1* as a critical negative regulator of stress-induced thrombopoiesis, functioning in part through intracellular calcium buffering and restriction of platelet-biased HSC expansion.

**Figure 2.**
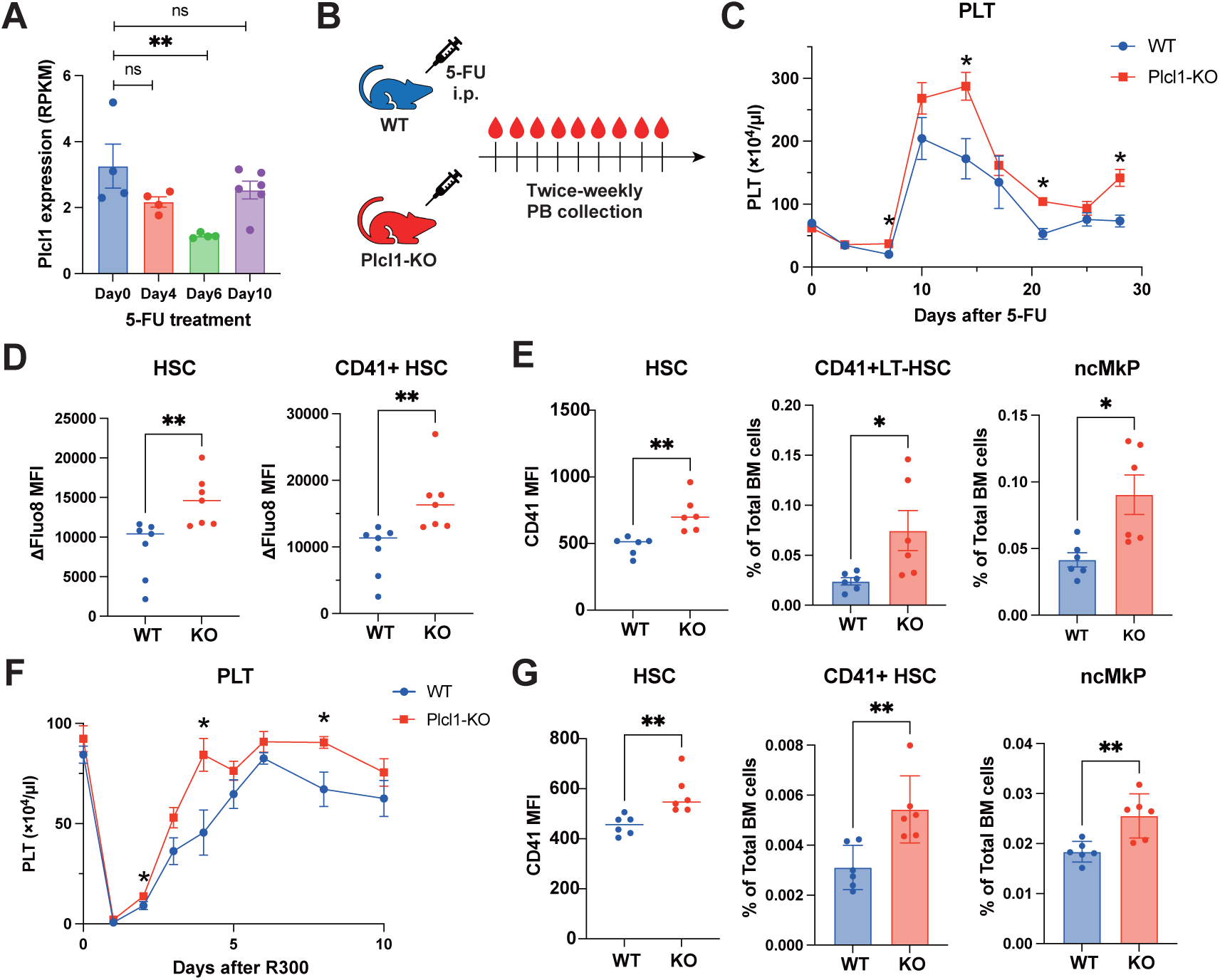
*Plcl1* deficiency accelerates platelet recovery under acute stress hematopoiesis. **(A)** Temporal dynamics of *Plcl1* expression in HSCs at 0, 4, 6, and 10 days following 5-FU administration, profiled using publicly available RNA-seq datasets (n = 4 for day 0, day 4, and day 6; n = 6 for day 10). **(B)** Schematic representation of the 5-FU-induced acute hematopoietic stress model used to assess peripheral blood recovery in WT and *Plcl1*-KO mice. **(C)** Peripheral blood platelet counts during recovery after 5-FU administration (150 mg/kg) in WT and *Plcl1*-KO mice (n = 6). **(D)** Intracellular Ca^2+^ concentrations in HSCs and CD41⁺ HSCs from WT and *Plcl1*-KO mice on day 6 post-5-FU, measured by Fluo-8 fluorescence intensity (n = 5). **p < 0.01. **(E)** CD41 mean fluorescence intensity (MFI) in HSCs and frequencies of CD41⁺ HSCs and non-canonical megakaryocyte progenitors (ncMkPs) in WT and *Plcl1*-KO mice on day 12 post-5-FU (n = 6). **(F)** Temporal analysis of peripheral platelet reconstitution following anti-CD42b-antibody-induced thrombocytopenia in WT and *Plcl1*-KO mice (n = 6). **(G)** Frequencies of CD41⁺ HSCs and ncMkPs in WT and *Plcl1*-KO mice on day 2 after anti-CD42b antibody administration (n = 5).

### *Plcl1* Deficiency Amplifies Age-Associated Platelet-Biased Hematopoiesis

Because aging and acute hematopoietic stress share common features, we examined the effects of *Plcl1* deficiency on age-associated HSC behavior. Analysis of public transcriptomic datasets revealed that *Plcl1* expression increases with age, alongside platelet-associated markers such as *Itga2b* (CD41), *Gp1ba* (CD42b), *Slamf1* (CD150), *Vwf*, *Itgb3* (CD61), and *Cd9* (Figure S4A)^40^, suggesting its potential role in age-associated hematopoietic remodeling. In aged *Plcl1*-KO mice, both the frequency and absolute number of HSCs were significantly elevated relative to those in aged WT controls, while ST-HSC, MPP2, and MPP3/4 compartments remained unchanged (**Figure 3A, Figure S4B**). This expansion of the HSC compartment was primarily driven by CD41⁺ HSCs, which exhibited a marked increase in both frequency and absolute number (**Figure 3A, Figure S4C**). Thus, aging and *Plcl1* deficiency cooperatively exacerbate megakaryocytic lineage bias. In line with these findings, aged *Plcl1*-KO mice exhibited a significant expansion of ncMkPs, while cMkPs remained unchanged (**Figure 3B, Figure S4D–E**), further supporting enhanced engagement of the bypass pathway with age in the absence of *Plcl1*. Functionally, competitive transplantation assays demonstrated that CD45.2⁺ donor cells from *Plcl1*-KO mice yielded significantly reduced peripheral blood chimerism across all lineages (**Figure 3C–G**). However, this defect was transient, as myeloid lineage reconstitution recovered to WT levels by four months post-transplantation (**Figure 3E**). These findings suggest that the observed impairment results from the dilution of functional HSCs within the expanded phenotypic pool, rather than from exhaustion or irreversible dysfunction. Analysis of lineage distribution further revealed a pronounced myeloid bias in donor-derived CD45.2⁺ cells, particularly at two months post-transplantation (**Figure 3H**), consistent with the aging-associated myeloid skewing of HSCs. Collectively, these results indicate that *Plcl1* deficiency accelerates key features of HSC aging, including HSC expansion, compromised reconstitution capacity, and myeloid-biased differentiation.

**Figure 3.**
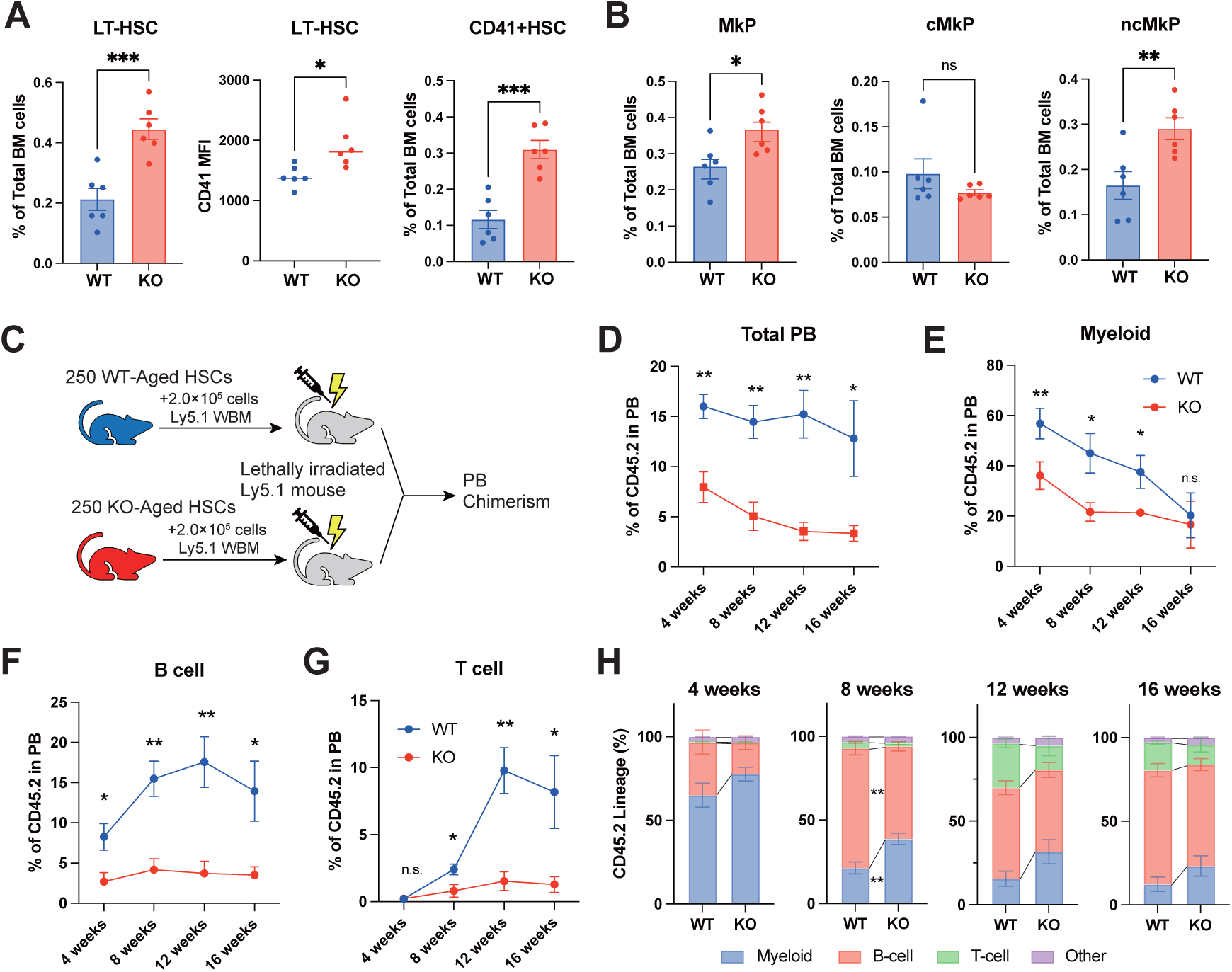
*Plcl1* mitigates HSC aging by suppressing the direct megakaryopoiesis bypass pathway. **(A)** Frequency of HSCs (% of total BM cells, left), CD41 mean fluorescence intensity (MFI) in HSCs (middle), and frequency of CD41⁺ HSCs (right) in aged WT and *Plcl1*-KO mice (n = 6). *p < 0.05; ***p < 0.001. **(B)** Frequencies of megakaryocyte progenitors (MkPs), canonical MkPs (cMkPs), and non-canonical MkPs (ncMkPs) as percentages of total BM cells in aged WT and *Plcl1*-KO mice (n = 6). *p < 0.05; **p < 0.01. **(C)** Schematic overview of competitive transplantation of aged WT or *Plcl1*-KO HSCs into young Ly5.1 recipient mice. **(D–G)** Donor-derived peripheral blood chimerism at 4, 8, 12, and 16 weeks post-transplantation in recipients of aged WT or *Plcl1*-KO HSCs (n = 5), showing chimerism in total peripheral blood cells (D), myeloid cells (E), B cells (F), and T cells (G). *p < 0.05; **p < 0.01; n.s., not significant. **(H)** Lineage distribution of donor-derived cells in peripheral blood at 4, 8, 12, and 16 weeks post-transplantation. *p < 0.05; **p < 0.01; not significant where not indicated.

### *Plcl1* Supports Calcium-Responsive Transcriptional Programs in Aged HSCs

To investigate the molecular basis of impaired calcium buffering in aged *Plcl1*-KO HSCs, we conducted RNA-seq-based transcriptomic analysis of HSCs from aged WT and *Plcl1*-KO mice (**Figure 4A–B**). Gene ontology analysis identified substantial downregulation of calcium-associated transcriptional programs, most notably the term “cellular response to calcium ion,” in *Plcl1*-KO HSCs (**Figure 4C–D**). The majority of these downregulated genes, including *Fos*, *Fosb*, *Jun*, *Junb*, *Nr4a1*, and *Egr1*, are members of the immediate-early gene (IEG) class^41^. These IEGs are well-established calcium-responsive transcriptional regulators^42^, indicating that *Plcl1* deficiency disrupts calcium-driven gene expression programs in aged HSCs (**Figure 4E**). Moreover, HSC-specific loss of *Junb*, *Nr4a1*, or *Egr1* has been reported to disrupt quiescence, promote aberrant cycling, expand the phenotypic HSC pool, and skew output toward myeloid lineages^43–45^. The reduced expression of these genes in *Plcl1*-KO HSCs may therefore contribute to aging-associated expansion and the preferential accumulation of CD41⁺ HSCs. To further dissect subset-specific transcriptional changes, we stratified aged HSCs into CD41⁻ and CD41⁺ subsets and conducted transcriptome profiling. Upon examination, CD41⁺ HSCs exhibited elevated expression of platelet-and proliferation-associated genes (*Itga2b, Itgb3, Vwf, Rrm1, Dnmt1*), consistent with their megakaryocytic priming and heightened cell cycle activity (**Figure S5A–E**)^37,46^. Importantly, calcium-signaling-related pathways remained suppressed in *Plcl1*-KO CD41⁺ HSCs, consistent with findings in bulk HSCs (**Figure 4E, Figure S5F**). Collectively, these results demonstrate that *Plcl1* is essential for proper transcriptional adaptation to calcium signaling in aged HSCs. Its absence diminishes HSC responsiveness to calcium-dependent cues, which exacerbates aging-associated expansion and megakaryocytic bias. Altogether, our findings establish *Plcl1* as a molecular brake that preserves HSC quiescence and lineage balance under age-related stress **(Figure 4F)**.

**Figure 4.**
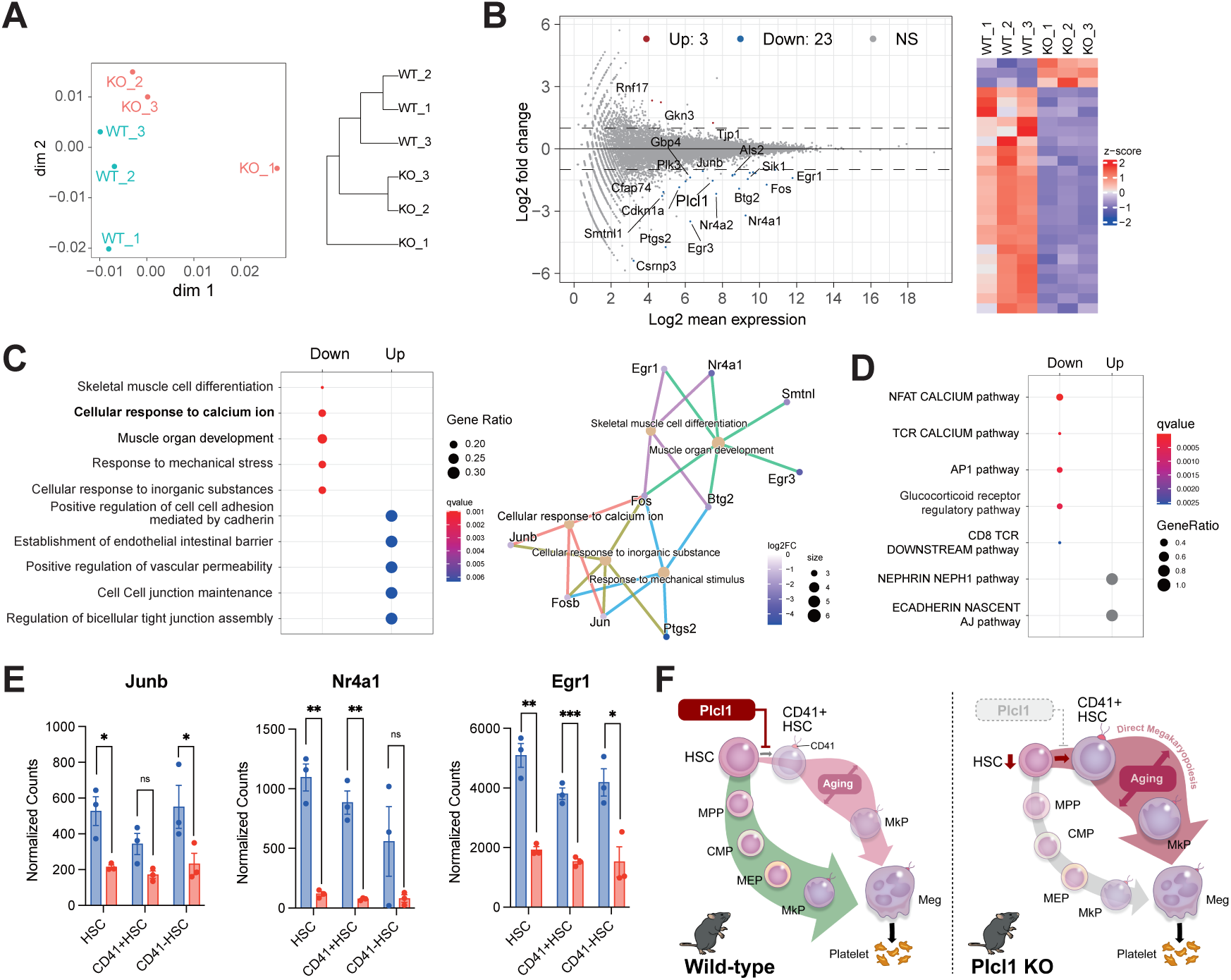
*Plcl1* safeguards calcium-dependent immediate early gene responses in aged HSCs. **(A)** Multidimensional scaling (MDS) and hierarchical clustering of transcriptomic profiles in HSCs from aged WT and *Plcl1*-KO mice (n = 3). **(B)** MA plot and hierarchical clustering heatmap illustrating differential gene expression (DEG) of aged HSCs from WT and *Plcl1*-KO mice (n = 3). **(C)** Enrichment analysis of GO biological processes based on DEGs between aged WT and *Plcl1*-KO HSCs, visualized using an over-representation bar plot and a gene–concept network (cnet) plot (n = 3). **(D)** Pathway interaction database enrichment analysis of DEGs from aged WT and *Plcl1*-KO HSCs (n = 3). **(E)** Normalized expression of calcium-responsive IEGs, namely *Junb*, *Nr4a1*, and *Egr1*, in aged HSCs from WT and *Plcl1*-KO mice (n = 3). n.s., not significant; *p < 0.05; **p < 0.01; ***p < 0.001. **(F)** Schematic model depicting how *Plcl1* deficiency in aged HSCs promotes platelet-biased HSC expansion and bypass megakaryopoiesis, resulting in altered hematopoiesis.

## Discussion

Extracellular calcium influx has been reported to regulate HSC behavior via L-type voltage-gated calcium channels, the calcium-sensing receptor, and purinergic receptors^10,47,48^. In contrast, the intracellular regulation of IP_3_-dependent Ca^2+^ release in HSCs remains unclear. Here, we demonstrate that *Plcl1*, a catalytically inactive IP_3_-binding protein, is enriched in the most quiescent HSC subset and modulates intracellular Ca^2+^ dynamics. Functionally, loss of *Plcl1* increased CD41⁺-megakaryocyte-primed HSCs at steady state, exaggerated platelet rebound during acute stress, and intensified aging-associated HSC expansion, and myeloid skewing, followed by diminished activation of Ca^2+^-responsive transcriptional programs. Together, these findings suggest that *Plcl1* acts as an intracellular modulator of Ca^2+^ signaling that helps restrain premature megakaryocytic priming during stress and aging.

Under steady-state conditions, *Plcl1* deficiency resulted in diminished basal Ca^2+^ levels in LT-HSCs. Consistent with this, previous biochemical studies indicate that *Plcl1* binds IP_3_ with high affinity, prolonging its half-life by competing with IP_3_ 5-phosphatase^26,49,50^. This binding is thought to sustain localized IP_3_–IP_3_R interactions and support basal Ca^2+^ release from the endoplasmic reticulum (ER), thereby enabling Ca^2+^-dependent transcriptional programs in quiescent HSCs. In contrast, following 5-FU treatment, intracellular Ca^2+^ levels in HSCs typically surge around day 6 as part of the regenerative rebound^10^, and in our study this Ca^2+^ surge was further amplified in *Plcl1*-KO HSCs. These findings suggest that *Plcl1* attenuates stress-induced Ca^2+^ release by sequestering IP_3_ and limiting excessive propagation of IP_3_R-mediated signaling^26,51^. Furthermore, analogous modulatory roles for *Plcl1* in IP_3_-mediated Ca^2+^ signaling have been documented in other non-hematopoietic contexts^26,28^. These support the view that *Plcl1* functions not as a canonical signaling effector but rather as a context-dependent modulator that restrains excessive Ca^2+^ transients and prevents premature or maladaptive lineage commitment. Thus, *Plcl1* acts primarily to modulate Ca^2+^ signaling dynamics rather than fundamentally altering core transcriptional networks and may influence the platelet-biased output of HSCs.

During physiological aging, cytosolic and mitochondrial Ca^2+^ levels progressively increase in HSCs, promoting a gradual loss of quiescence and accelerating HSC aging^14,52^. A key mechanism driving this increase involves elevated CD38 activity in aged HSCs, which generates metabolites such as cADPR that stimulate ER-based Ca^2+^ release, primarily through ryanodine receptors^52^. Our findings suggest that the pronounced aging-associated phenotypes in aged and *Plcl1*-KO HSCs likely result from this synergistic disruption of Ca²⁺ homeostasis. Notably, although *Plcl1* deletion reduces basal cytosolic Ca^2+^ levels in young HSCs, this effect was not observed in aged HSCs. This phenomenon is presumably due to the dominant influence of aging-associated intracellular Ca^2+^ influx^52^. Furthermore, aged *Plcl1*-KO HSCs display reduced expression of Ca^2+^-responsive IEGs, including *JunB*, *Egr1*, and *Nr4a1*, which are critical for maintaining quiescence and enabling appropriate stress responses^43,45,53^. Because IEG activation depends not on global Ca^2+^ elevation but on the finely tuned dynamics of Ca^2+^ spikes, including their amplitude, frequency, and duration,^53^ disruption of these parameters in aged *Plcl1*-KO HSCs likely decouples Ca^2+^ signaling from transcriptional output, thereby impairing adaptive transcriptional programs. Together, these findings identify *Plcl1* as a critical gatekeeper of Ca^2+^ dynamics that preserves Ca^2+^-dependent transcriptional responses in aged HSCs, thereby maintaining stem cell stability under aging-associated stress.

This study has several limitations. We were unable to directly assess IP_3_ buffering or Ca^²⁺^ oscillatory dynamics in primary HSCs. Moreover, potential redundancy with other regulators such as *Plcl2* or IP_3_Rs remains unresolved. Analysis of a publicly available human hematopoiesis dataset (GSE42519) indicated that *PLCL1* transcript levels in human HSCs (Lin⁻ CD34⁺ CD38⁻ CD90⁺ CD45RA⁻) were approximately 1.5-fold higher than in MPPs and granulocyte/macrophage progenitors and 1.7-fold higher than in common myeloid progenitors and megakaryocyte/erythrocyte progenitors. These data suggest that *Plcl1* may also be enriched in human HSCs at the transcript level. However, *Plcl1* expression at the protein level, as well as its functional contribution to human HSC regulation, remains to be validated in future studies.

In summary, our findings identify *Plcl1* as a critical modulator of Ca^2+^ signaling that preserves HSC quiescence and mitigates age- and stress-associated megakaryocyte priming. These insights establish a mechanistic link between Ca^2+^ homeostasis and stem cell fate and highlight *Plcl1* as a potential target for sustaining balanced hematopoiesis during aging and following therapeutic intervention.

## Acknowledgments

We thank Dr. Masato Hirata (Fukuoka Dental College) for generously providing the Plcl1 knockout mice. We are also grateful to Takako Ideue, Rina Iwata, Miho Kataoka, and Shiori Shikata for their excellent technical assistance. We appreciate the support of the core facilities of the International Research Center for Medical Sciences (IRCMS), Kumamoto University, for technical support, and the Center for Animal Resources and Development (CARD), Kumamoto University, for excellent animal care. We also thank the FACS Core and the Mouse Core at the Institute of Medical Science, the University of Tokyo. This work was supported by the Japan Society for the Promotion of Science (JSPS) Grant-in-Aid for Young Scientists (TY), a JSPS Grant-in-Aid for Young Scientists (TY, 24K19227), a JSPS Grant-in-Aid for Challenging Explanatory Research (YT, 24K11520), a JSPS Grant-in-Aid for (21K19500), a grant from SENSHIN Medical Research Foundation (TY), a Grant-in-Aid for Scientific research from MOCHIDA Memorial Foundation (TY and YT), the Japanese Society of Hematology Research grant (TY), and Kanehara Ichiro Award (TS). This research was also supported by MEXT Promotion of Distinctive Joint Usage/Research Center Support Program Grant Numbers JPMXP0724020288 at the Advanced Medical Research Center, Yokohama City University.

## Authorship

T.Y. and Y.T. conceived the project, designed and performed most of the experiments, analyzed and interpreted the data, and wrote the manuscript. T.F., T.I., R.I., M.K., S.S., K.W., K.K., D.K., Y.A., T.U., assisted with the experiments. T.F., H.T., S.G., T.K., A.N., advised on the data interpretation, discussed and made suggestions for this study. T.S. supervised the project, interpreted the data, and participated in writing the manuscript.

---

**Supplementary Figure 1.**
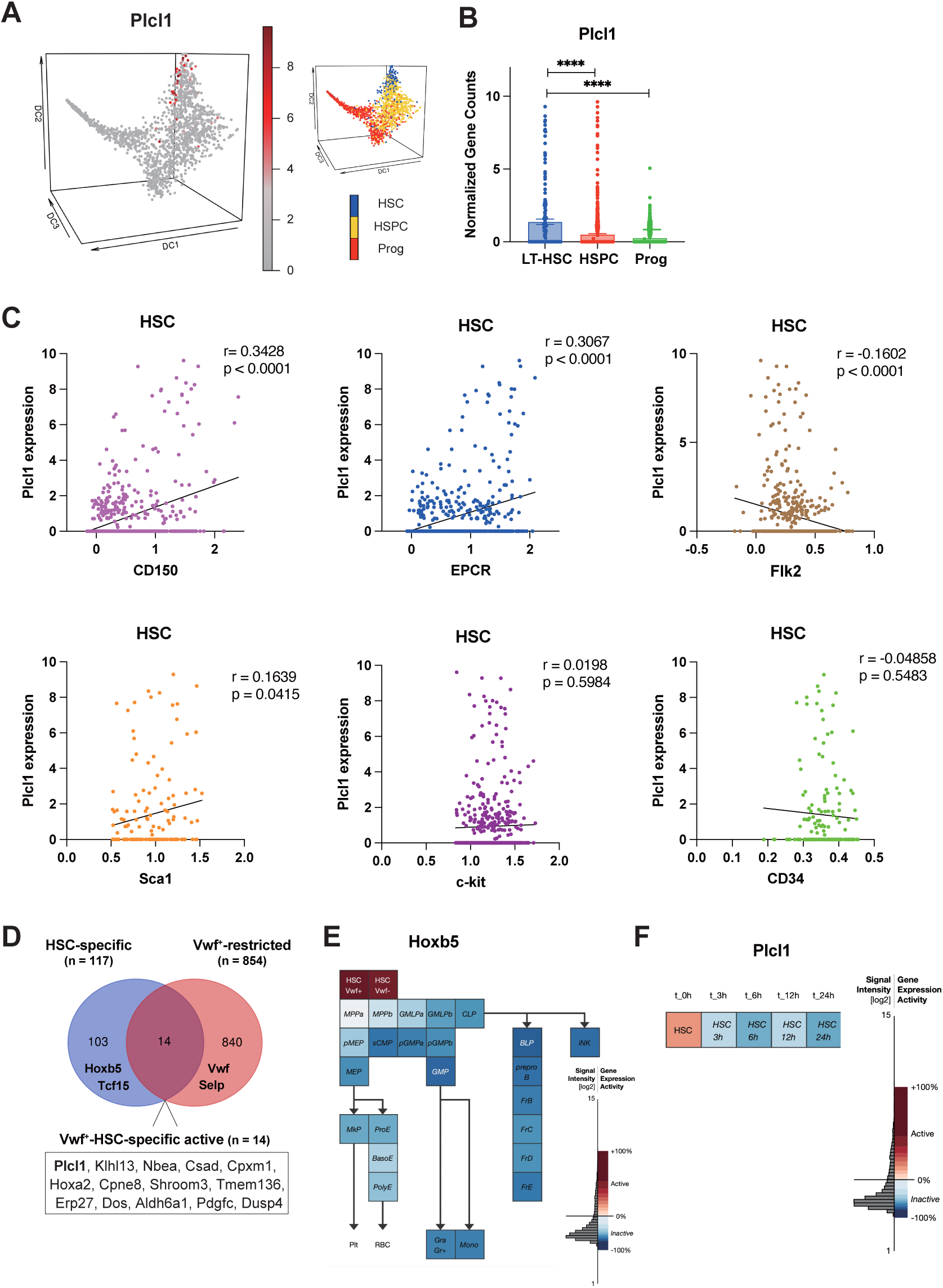
*Plcl1* marks quiescent HSCs in the G₀ phase and associates with elevated intracellular calcium levels. (**A–C**) Single-cell RNA-seq analysis from the Single-Cell Gene Expression Atlas database. **(A)** PCA plot depicting enrichment of *Plcl1* expression (red) in HSCs. **(B)** Comparison of *Plcl1* expression across gated populations: Prog (Lin⁻ Sca-1⁻ c-Kit⁺), HSPCs (Lin⁻ Sca-1⁺ c-Kit⁺), and HSCs (Lin⁻ Sca-1⁺ c-Kit⁺ CD34⁻ Flk2⁻). ****p < 0.0001. **(C)** Correlative analysis of *Plcl1* expression with cell surface antigens (CD150, EPCR, Flk2, Sca-1, c-Kit, and CD34) at the single-cell level. **(D)** Venn diagram from the Gene Expression Commons database showing that *Plcl1* is among the Vwf⁺-HSC-specific active genes shared between HSC-specific genes (n = 117) and Vwf⁺-restricted genes (n = 854). **(E)** Summary from Gene Expression Commons illustrating an example of an HSC-specific gene, with *Hoxb5* displayed as a heatmap showing peak expression in HSCs. Cell population abbreviations: MPP, multipotent progenitor; GMP, granulocyte/macrophage progenitor; MEP, megakaryocyte/erythrocyte progenitor; CMP, common myeloid progenitor; MkP, megakaryocyte progenitor; CLP, common lymphoid progenitor; BasoE, basophilic erythroblast; PolyE, polyorthochromatic erythroblast; Mono, monocyte; Gra, granulocyte; BLP, B-lymphoid progenitor; preproB, pre-pro B cell; MzB, marginal zone B cell; FoB, follicular B cell; iNK, intermediate natural killer cell; mNK, mature natural killer cell. For further details, see Chen et al. (Nature 530, 223–227, 2016). **(F)** Bar graph depicting rapid downregulation of *Plcl1* expression in HSCs after ex vivo culture for 3, 6, 12, and 24 h.

**Supplementary Figure 2.**
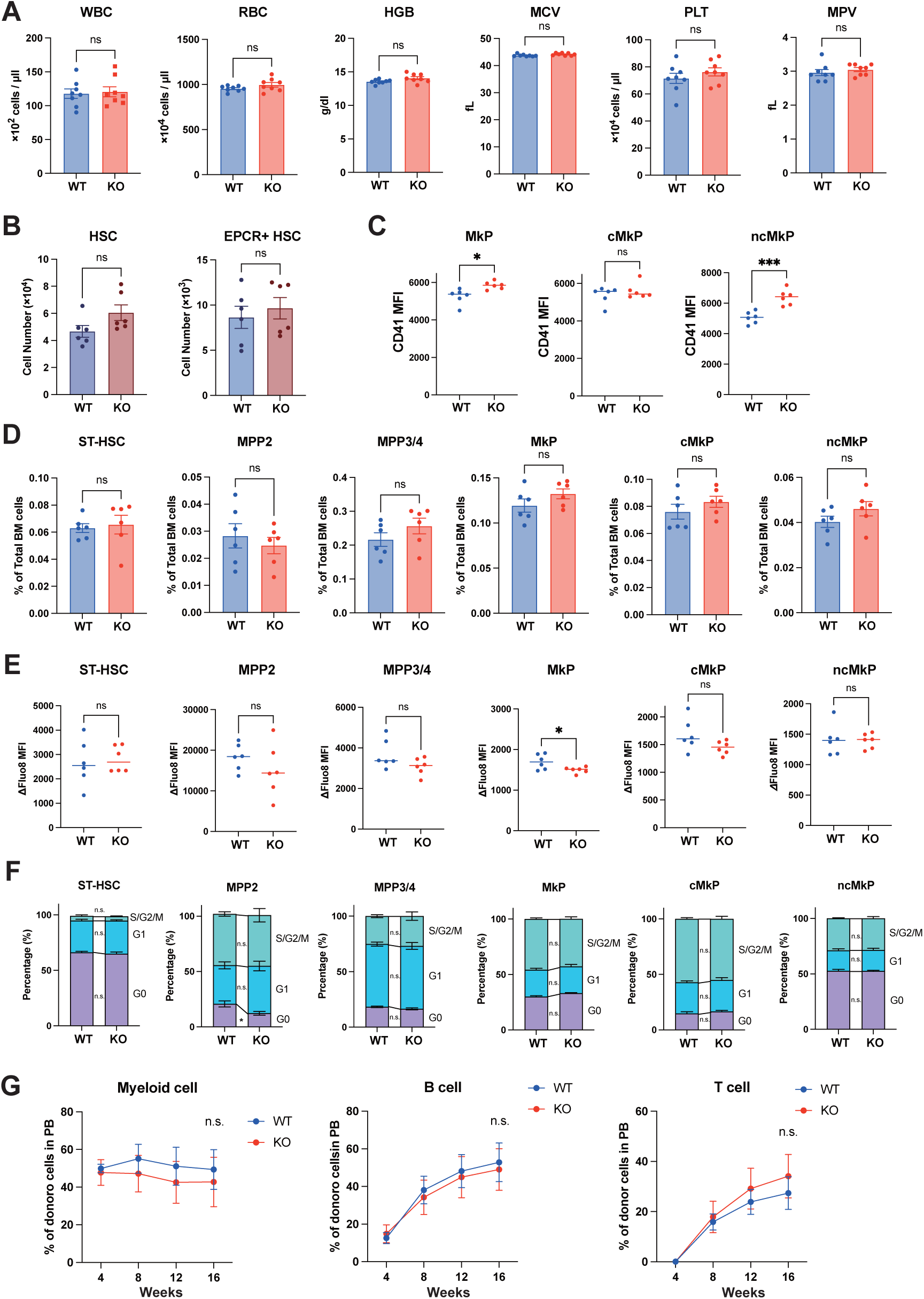

**Supplementary Figure 3.**
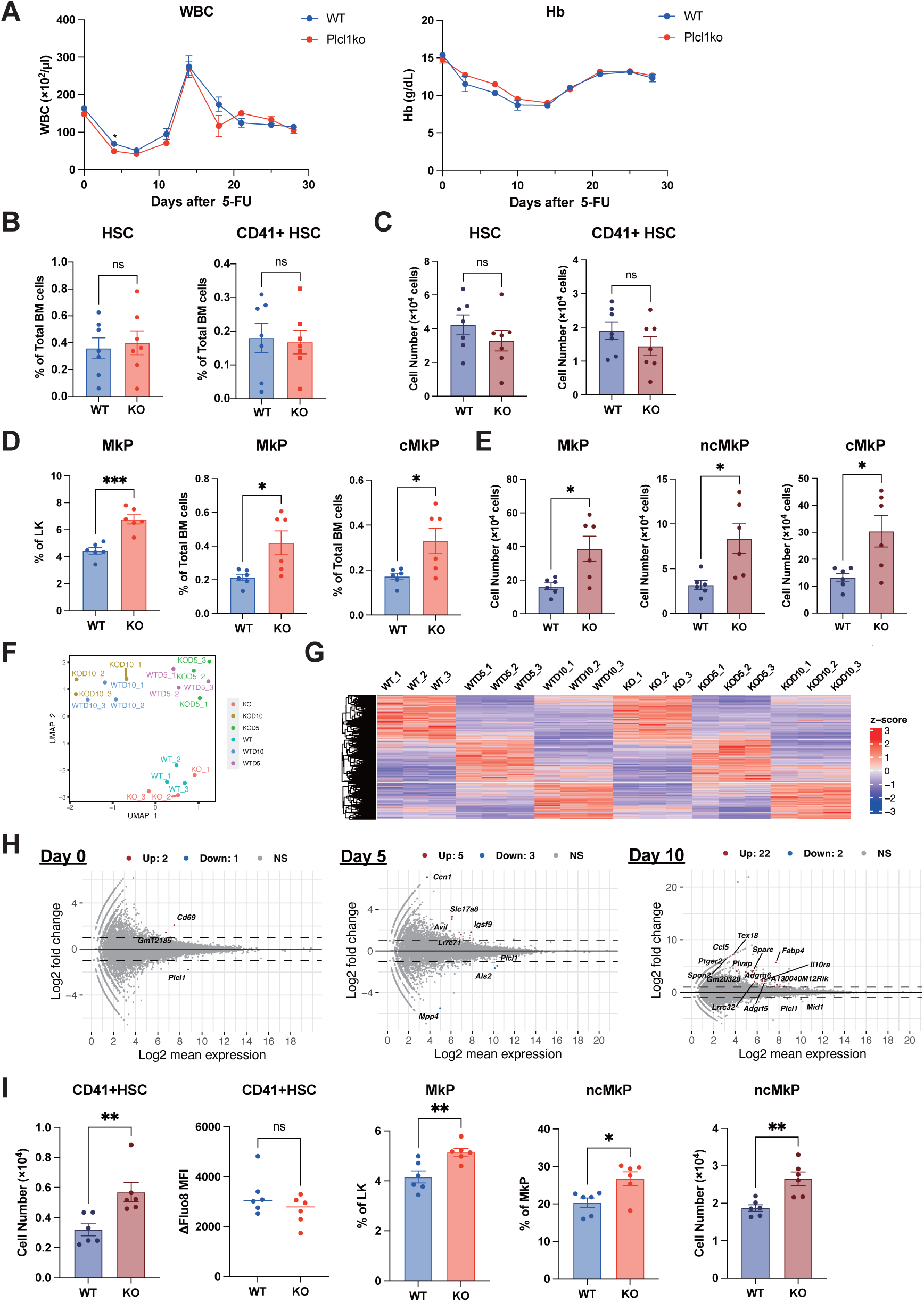
Loss of *Plcl1* selectively enhances platelet-biased responses without affecting global hematopoiesis under acute hematopoietic stress. (**A**) Peripheral blood white blood cell (WBC) counts (left) and hemoglobin (HGB) levels (right) during hematopoietic recovery following 5-FU (150 mg/kg) administration in WT and *Plcl1*-KO mice (n = 6). (**B** and **C**) Frequencies (**B**) and absolute numbers (**C**) of HSCs (L-ESLAM; Lineage⁻ EPCR⁺ CD150⁺ CD48⁻) and CD41⁺ HSCs at day 6 after 5-FU administration in WT and *Plcl1*-KO mice (n = 6). ns: not significant. (**D** and **E**) Frequencies (**D**) and absolute numbers (**E**) of MkPs at day 12 after 5-FU administration in WT and *Plcl1*-KO mice (n = 6). \**p* < 0.05; \*\*\**p* < 0.001. (**F** and **G**) UMAP projection (**F**) and unsupervised clustering (**G**) of transcriptomic profiles obtained by RNA-seq from HSCs isolated at days 0, 5, and 10 following 5-FU administration in WT and *Plcl1*-KO mice (n = 3). (**H**) MA plot displaying DEGs between WT and *Plcl1*-KO L-ESLAM cells at days 0, 5, and 10 after 5-FU treatment based on RNA-seq data. (**I**) Hematopoietic response at day 2 after administration of anti-CD42b antibody (R300) in WT and *Plcl1*-KO mice. The absolute number of CD41⁺ HSCs, intracellular calcium levels (ΔFluo-8 MFI) in CD41⁺ HSCs, MkP and ncMkP frequencies (as a percentage of MkPs), and absolute number of ncMkPs (n = 6) are indicated. ns: not significant; \**p* < 0.05; \*\**p* < 0.01; \*\*\**p* < 0.001; \*\*\*\**p* < 0.0001.

**Supplementary Figure 4.**
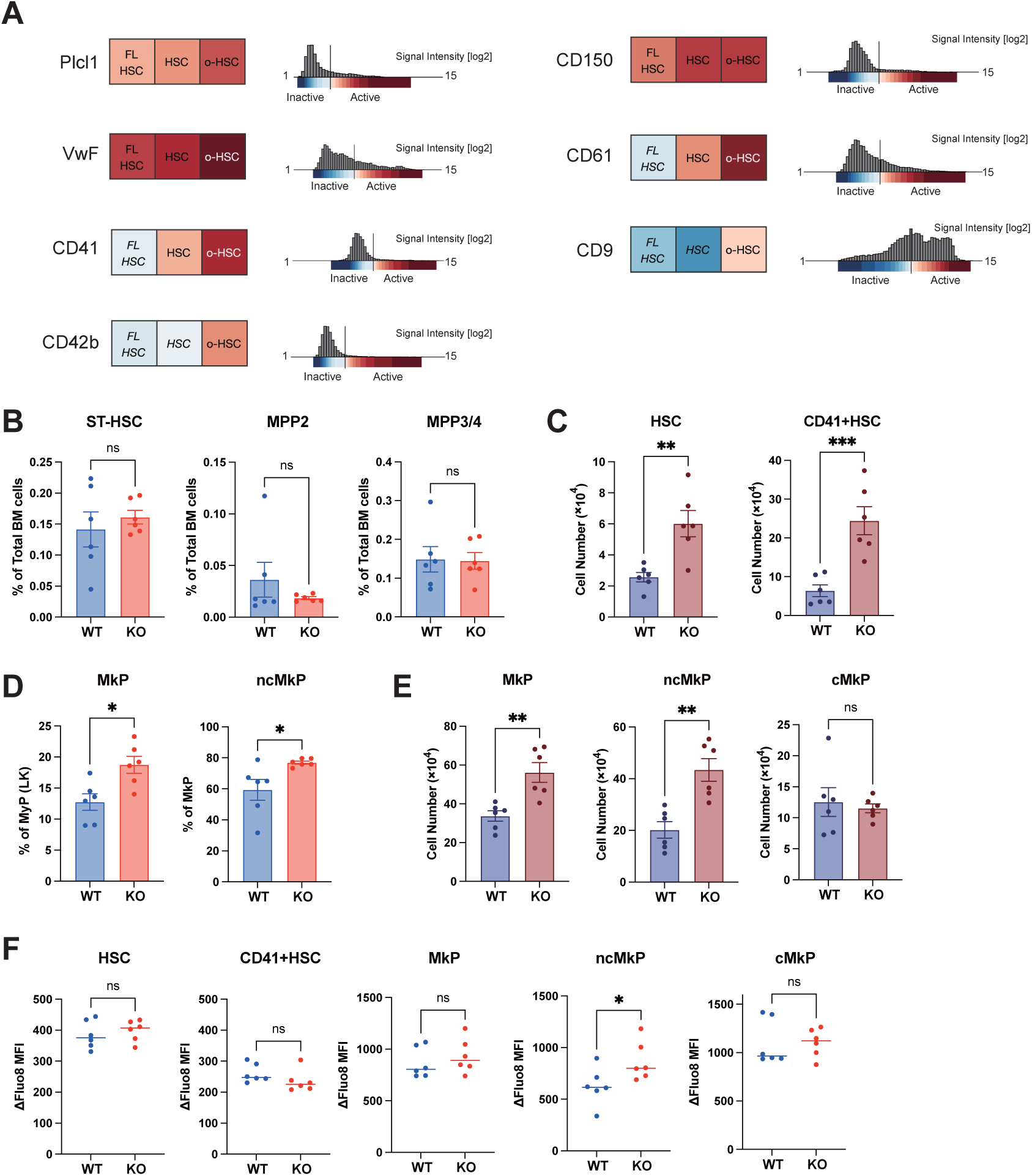
*Plcl1* deficiency exacerbates age-associated megakaryocyte bias and alters HSC composition. **(A)** Gene expression analysis using the Gene Expression Commons platform (Seita et al., PLoS One, 2012) indicates that *Plcl1* expression increases with age, in parallel with other platelet-biased HSC markers including *Itga2b* (CD41), *Gp1ba* (CD42b), *Cd150*, *Vwf*, *Cd9*, *Itgb3* (CD61), and *Gabrr1*. **(B)** Frequencies of KSL subfractions, namely ST-HSCs, MPP2, and MPP3/4, expressed as percentages of total bone marrow cells (n = 6). ns, not significant. **(C)** Absolute numbers of HSCs and CD41⁺ HSCs in aged WT and *Plcl1*-KO mice (n = 6). \**p* < 0.05; \*\**p* < 0.01; \*\*\**p* < 0.001; ns, not significant. **(D)** Frequencies of MkPs (% of LK) and ncMkPs (% of MkPs) in aged WT and *Plcl1*-KO mice. \**p* < 0.05. **(E)** Absolute numbers of MkPs, ncMkPs, and cMkPs in aged WT and *Plcl1*-KO mice (n = 6). \**p* < 0.05; \*\**p* < 0.01; ns, not significant. **(F)** Intracellular calcium levels (ΔFluo-8 mean fluorescence intensity) in HSCs, CD41⁺ HSCs, MkPs, ncMkPs, and cMkPs in aged WT and *Plcl1*-KO mice (n = 6). ns, not significant. \**p* < 0.05.

**Supplementary Figure 5.**
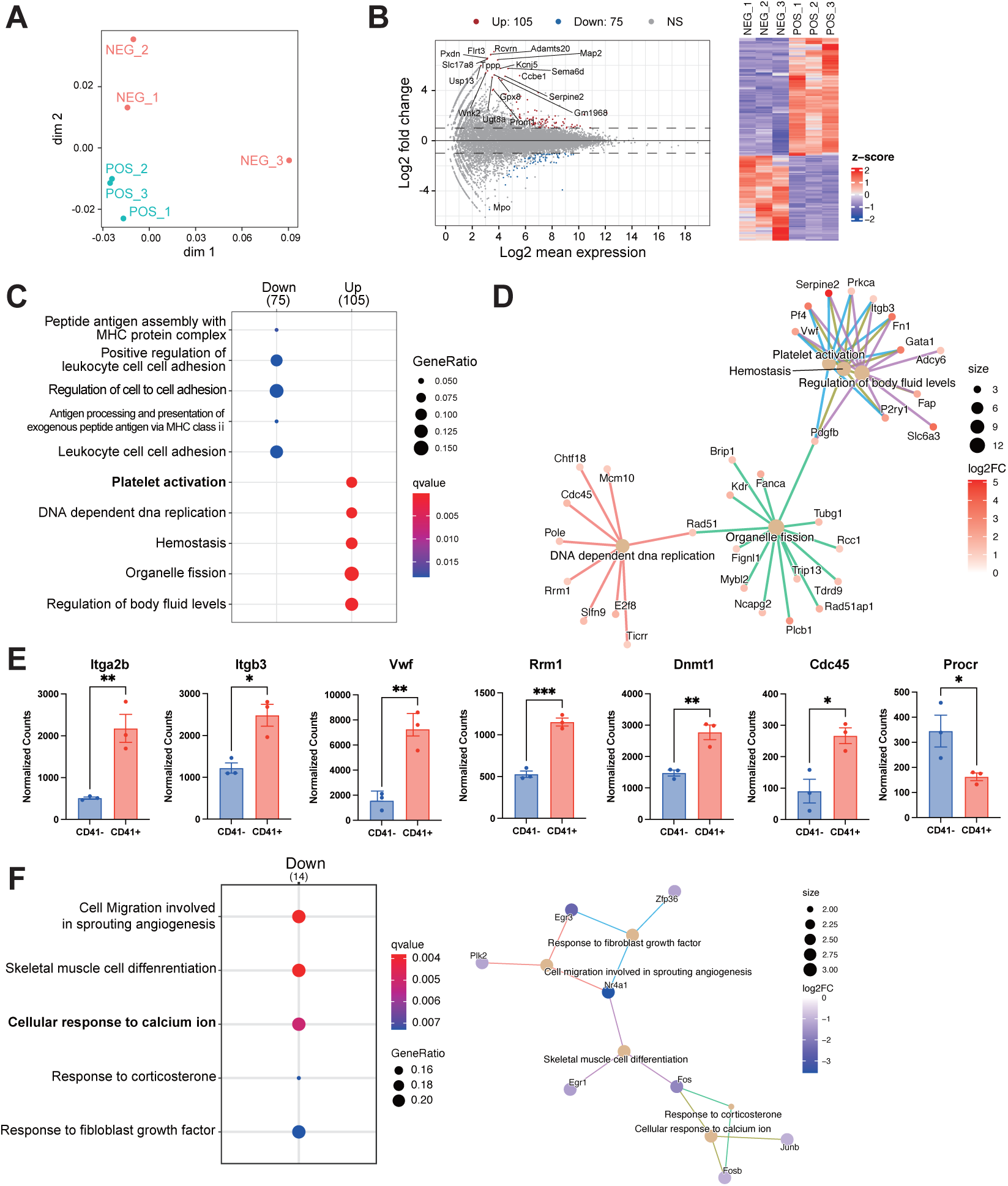
Transcriptomic profiling of aged CD41⁺ HSCs and *Plcl1*-deficient CD41⁺ HSCs. **(A)** Transcriptomic profiles of aged CD41⁻ and CD41⁺ WT HSCs analyzed via RNA-seq and visualized using multi-dimensional scaling (MDS) and hierarchical clustering (n = 3). **(B)** MA plot and hierarchical clustering heatmap depicting results of DEG analysis comparing aged CD41⁻ and CD41⁺ WT HSCs (n = 3). (**C, D**) GO biological process enrichment analysis of DEGs between aged CD41⁻ and CD41⁺ HSCs, visualized as an over-representation bar plot (**C**) and a gene–concept network (cnet) plot **(D)** (n = 3). **(E)** Normalized expression levels of surface markers associated with platelet-biased HSCs (*Itga2b*, *Itgb3*, *Vwf*) and cell cycle–related genes (*Rrm1*, *Dnmt1*, *Cdc45*, *Procr*) in aged CD41⁺ and CD41⁻ HSCs (n = 3). \**p* < 0.05; \*\**p* < 0.01; \*\*\**p* < 0.001. **(F)** GO biological process enrichment analysis of DEGs between aged CD41⁺ WT and *Plcl1*-KO HSCs, visualized as an over-representation bar plot (n = 3).

## References

1. Wilson A, Laurenti E, Oser G, et al. Hematopoietic stem cells reversibly switch from dormancy to self-renewal during homeostasis and repair. Cell. 2008;135(6):1118–1129.

2. Foudi A, Hochedlinger K, Van Buren D, et al. Analysis of histone 2B-GFP retention reveals slowly cycling hematopoietic stem cells. Nat Biotechnol. 2009;27(1):84–90.

3. Yamazaki S, Iwama A, Takayanagi S, et al. TGF-beta as a candidate bone marrow niche signal to induce hematopoietic stem cell hibernation. Blood. 2009;113(6):1250–1256.

4. Tothova Z, Kollipara R, Huntly BJ, et al. FoxOs are critical mediators of hematopoietic stem cell resistance to physiologic oxidative stress. Cell. 2007;128(2):325–339.

5. Takubo K, Goda N, Yamada W, et al. Regulation of the HIF-1alpha level is essential for hematopoietic stem cells. Cell Stem Cell. 2010;7(3):391–402.

6. Matsumoto A, Takeishi S, Kanie T, et al. p57 is required for quiescence and maintenance of adult hematopoietic stem cells. Cell Stem Cell. 2011;9(3):262–271.

7. Hock H, Hamblen MJ, Rooke HM, et al. Gfi-1 restricts proliferation and preserves functional integrity of haematopoietic stem cells. Nature. 2004;431(7011):1002–1007.

8. Morita Y, Ema H, Nakauchi H. Heterogeneity and hierarchy within the most primitive hematopoietic stem cell compartment. J Exp Med. 2010;207(6):1173–1182.

9. Dykstra B, Kent D, Bowie M, et al. Long-term propagation of distinct hematopoietic differentiation programs in vivo. Cell Stem Cell. 2007;1(2):218–229.

10. Umemoto T, Hashimoto M, Matsumura T, et al. Ca(2+)-mitochondria axis drives cell division in hematopoietic stem cells. J Exp Med. 2018;215(8):2097–2113.

11. Luchsinger LL, de Almeida MJ, Corrigan DJ, et al. Mitofusin 2 maintains haematopoietic stem cells with extensive lymphoid potential. Nature. 2016;529(7587):528–531.

12. Luchsinger LL, Strikoudis A, Danzl NM, et al. Harnessing Hematopoietic Stem Cell Low Intracellular Calcium Improves Their Maintenance In Vitro. Cell Stem Cell. 2019;25(2):225–240.e227.

13. Fukushima T, Tanaka Y, Hamey FK, et al. Discrimination of Dormant and Active Hematopoietic Stem Cells by G(0) Marker Reveals Dormancy Regulation by Cytoplasmic Calcium. Cell Rep. 2019;29(12):4144–4158.e4147.

14. Song Z, Park SH, Mu WC, et al. An NAD(+)-dependent metabolic checkpoint regulates hematopoietic stem cell activation and aging. Nat Aging. 2024;4(10):1384–1393.

15. Poscablo DM, Worthington AK, Smith-Berdan S, et al. An age-progressive platelet differentiation path from hematopoietic stem cells causes exacerbated thrombosis. Cell. 2024;187(12):3090–3107.e3021.

16. Rodriguez-Fraticelli AE, Wolock SL, Weinreb CS, et al. Clonal analysis of lineage fate in native haematopoiesis. Nature. 2018;553(7687):212–216.

17. Sanjuan-Pla A, Macaulay IC, Jensen CT, et al. Platelet-biased stem cells reside at the apex of the haematopoietic stem-cell hierarchy. Nature. 2013;502(7470):232–236.

18. Kowalczyk MS, Tirosh I, Heckl D, et al. Single-cell RNA-seq reveals changes in cell cycle and differentiation programs upon aging of hematopoietic stem cells. Genome Res. 2015;25(12):1860–1872.

19. Morcos MNF, Li C, Munz CM, et al. Fate mapping of hematopoietic stem cells reveals two pathways of native thrombopoiesis. Nat Commun. 2022;13(1):4504.

20. Carrelha J, Mazzi S, Winroth A, et al. Alternative platelet differentiation pathways initiated by nonhierarchically related hematopoietic stem cells. Nat Immunol. 2024;25(6):1007–1019.

21. Li JJ, Liu J, Li YE, et al. Differentiation route determines the functional outputs of adult megakaryopoiesis. Immunity. 2024;57(3):478–494.e476.

22. Haas S, Hansson J, Klimmeck D, et al. Inflammation-Induced Emergency Megakaryopoiesis Driven by Hematopoietic Stem Cell-like Megakaryocyte Progenitors. Cell Stem Cell. 2015;17(4):422–434.

23. Walter D, Lier A, Geiselhart A, et al. Exit from dormancy provokes DNA-damage-induced attrition in haematopoietic stem cells. Nature. 2015;520(7548):549–552.

24. Hirata M, Kukita M, Sasaguri T, et al. Increase in Ca2+ permeability of intracellular Ca2+ store membrane of saponin-treated guinea pig peritoneal macrophages by inositol 1,4,5-trisphosphate. J Biochem. 1985;97(6):1575–1582.

25. Hirata M, Sasaguri T, Hamachi T, et al. Irreversible inhibition of Ca2+ release in saponin-treated macrophages by the photoaffinity derivative of inositol-1, 4, 5-trisphosphate. Nature. 1985;317(6039):723–725.

26. Harada K, Takeuchi H, Oike M, et al. Role of PRIP-1, a novel Ins(1,4,5)P3 binding protein, in Ins(1,4,5)P3-mediated Ca2+ signaling. J Cell Physiol. 2005;202(2):422–433.

27. Muter J, Brighton PJ, Lucas ES, et al. Progesterone-Dependent Induction of Phospholipase C-Related Catalytically Inactive Protein 1 (PRIP-1) in Decidualizing Human Endometrial Stromal Cells. Endocrinology. 2016;157(7):2883–2893.

28. Murakami A, Matsuda M, Harada Y, et al. Phospholipase C-related, but catalytically inactive protein (PRIP) up-regulates osteoclast differentiation via calcium-calcineurin-NFATc1 signaling. J Biol Chem. 2017;292(19):7994–8006.

29. Kanematsu T, Jang IS, Yamaguchi T, et al. Role of the PLC-related, catalytically inactive protein p130 in GABA(A) receptor function. Embo j. 2002;21(5):1004–1011.

30. Umemoto T, Johansson A, Ahmad SAI, et al. ATP citrate lyase controls hematopoietic stem cell fate and supports bone marrow regeneration. Embo j. 2022;41(8):e109463.

31. Kubota S, Sun Y, Morii M, et al. Chromatin modifier Hmga2 promotes adult hematopoietic stem cell function and blood regeneration in stress conditions. Embo j. 2024;43(13):2661–2684.

32. Etoh K, Nakao M. A web-based integrative transcriptome analysis, RNAseqChef, uncovers the cell/tissue type-dependent action of sulforaphane. J Biol Chem. 2023;299(6):104810.

33. Nestorowa S, Hamey FK, Pijuan Sala B, et al. A single-cell resolution map of mouse hematopoietic stem and progenitor cell differentiation. Blood. 2016;128(8):e20–31.

34. Seita J, Sahoo D, Rossi DJ, et al. Gene Expression Commons: an open platform for absolute gene expression profiling. PLoS One. 2012;7(7):e40321.

35. Yamamoto R, Morita Y, Ooehara J, et al. Clonal analysis unveils self-renewing lineage-restricted progenitors generated directly from hematopoietic stem cells. Cell. 2013;154(5):1112–1126.

36. Psaila B, Wang G, Rodriguez-Meira A, et al. Single-Cell Analyses Reveal Megakaryocyte-Biased Hematopoiesis in Myelofibrosis and Identify Mutant Clone-Specific Targets. Mol Cell. 2020;78(3):477–492.e478.

37. Kristiansen TA, Zhang Q, Vergani S, et al. Developmental cues license megakaryocyte priming in murine hematopoietic stem cells. Blood Adv. 2022;6(24):6228–6241.

38. Chenaille PJ, Steward SA, Ashmun RA, et al. Prolonged thrombocytosis in mice after 5-fluorouracil results from failure to down-regulate megakaryocyte concentration. An experimental model that dissociates regulation of megakaryocyte size and DNA content from megakaryocyte concentration. Blood. 1990;76(3):508–515.

39. Luis TC, Barkas N, Carrelha J, et al. Perivascular niche cells sense thrombocytopenia and activate hematopoietic stem cells in an IL-1 dependent manner. Nat Commun. 2023;14(1):6062.

40. Frisch BJ, Hoffman CM, Latchney SE, et al. Aged marrow macrophages expand platelet-biased hematopoietic stem cells via Interleukin1B. JCI Insight. 2019;5(10).

41. Fowler T, Sen R, Roy AL. Regulation of primary response genes. Mol Cell. 2011;44(3):348–360.

42. Ginty DD. Calcium regulation of gene expression: isn’t that spatial? Neuron. 1997;18(2):183–186.

43. Freire PR, Conneely OM. NR4A1 and NR4A3 restrict HSC proliferation via reciprocal regulation of C/EBPα and inflammatory signaling. Blood. 2018;131(10):1081–1093.

44. Santaguida M, Schepers K, King B, et al. JunB protects against myeloid malignancies by limiting hematopoietic stem cell proliferation and differentiation without affecting self-renewal. Cancer Cell. 2009;15(4):341–352.

45. Min IM, Pietramaggiori G, Kim FS, et al. The transcription factor EGR1 controls both the proliferation and localization of hematopoietic stem cells. Cell Stem Cell. 2008;2(4):380–391.

46. Ramasz B, Krüger A, Reinhardt J, et al. Hematopoietic stem cell response to acute thrombocytopenia requires signaling through distinct receptor tyrosine kinases. Blood. 2019;134(13):1046–1058.

47. Adams GB, Chabner KT, Alley IR, et al. Stem cell engraftment at the endosteal niche is specified by the calcium-sensing receptor. Nature. 2006;439(7076):599–603.

48. Wang N, Yin J, You N, et al. TWIST1 preserves hematopoietic stem cell function via the CACNA1B/Ca2+/mitochondria axis. Blood. 2021;137(21):2907–2919.

49. Jefferson AB, Majerus PW. Properties of type II inositol polyphosphate 5-phosphatase. J Biol Chem. 1995;270(16):9370–9377.

50. De Smedt F, Boom A, Pesesse X, et al. Post-translational modification of human brain type I inositol-1,4,5-trisphosphate 5-phosphatase by farnesylation. J Biol Chem. 1996;271(17):10419–10424.

51. Takeuchi H, Oike M, Paterson HF, et al. Inhibition of Ca(2+) signalling by p130, a phospholipase-C-related catalytically inactive protein: critical role of the p130 pleckstrin homology domain. Biochem J. 2000;349(Pt 1):357–368.

52. Jankowski CSR, Weichhart T. CD38 and the mitochondrial calcium uniporter contribute to age-related hematopoietic stem cell dysfunction. Immunometabolism (Cobham). 2024;6(4):e00048.

53. Berridge MJ, Bootman MD, Roderick HL. Calcium signalling: dynamics, homeostasis and remodelling. Nat Rev Mol Cell Biol. 2003;4(7):517–529.

54. Oki T, Nishimura K, Kitaura J, et al. A novel cell-cycle-indicator, mVenus-p27K-, identifies quiescent cells and visualizes G0-G1 transition. Sci Rep. 2014;4:4012.

